# Cell intrinsic signaling in *MEN1* mutant pancreatic neuroendocrine tumors unveils novel signaling pathways associated with de-differentiation

**DOI:** 10.1101/2020.09.29.318873

**Authors:** Brenna A. Rheinheimer, Ronald L. Heimark, Adam D. Grant, Luis Camacho, Megha Padi, Tun Jie

## Abstract

**Objectives:** Preliminary genomic analysis of primary pancreatic neuroendocrine tumors revealed a complex mutational landscape with four common oncogenic events; however, critical activation pathways responsible for pancreatic neuroendocrine tumor progression and metastasis have yet to be elucidated. Here, we analyzed six primary pancreatic neuroendocrine tumors to determine which pathways are deregulated and responsible for progression.

**Methods:** Selected genomic profiling of six primary pancreatic neuroendocrine tumors was performed using the Ion Torrent Comprehensive Cancer Panel with matched transcriptomes analyzed by Affymetrix Clariom D arrays. Validation of gene expression changes were measured by quantitative PCR using TaqMan assays and immunohistochemistry on tumor specimens.

**Results:** *MEN1* was mutated in half (50%) of our sequenced tumors while *FGFR3* was mutated in 2/6 (33%). Transcriptome analysis revealed that *ITGA2* and *EZH2* were overexpressed in *MEN1* mutant tumors whereas *ALK* and *VEGFA* were overexpressed in *FGFR3* mutant tumors. Immunohistochemistry revealed increased nuclear ITGA2 and EZH2 staining along with increased VE-Cadherin staining and loss of membranous E-cadherin localization in *MEN1* and *FGFR3* mutant tumors.

**Conclusions:** Our results suggest that pancreatic neuroendocrine tumors containing *MEN1* and *FGFR3* mutations are more aggressive and de-differentiated than their wild-type counterparts. Additionally, we provide novel chemotherapeutic target FGFR3 for patients with this disease.

## Introduction

Pancreatic neuroendocrine tumors (PanNETs), while rare, are one of the fastest growing cancer diagnoses due to improved detection of asymptomatic malignancies. While these tumors are thought to develop from cells within the islets of Langerhans, the initiating cell type has yet to be identified. Only 10% of PanNETs are functional and related to a clinical syndrome associated with hypersecretion of hormones. The other 90% of PanNETs are non-functional and develop asymptomatically leading to delayed clinical presentation^1^. Therefore, the majority of PanNET patients present with locally advanced (21%) or metastatic (60%) disease at diagnosis^2^. The overall five-year survival rate of patients with localized non-functional PanNETs is approximately 31% which drops to 18% in patients with metastatic disease^3^. Currently, the only curative treatment option for PanNETs is surgical resection; however most tumors are diagnosed beyond the criteria of resectability. Therefore, treatment of PanNETs is largely empirically based from a lack of scientific evidence that supports how and when to treat patients with aggressive or metastatic disease.

Multiple endocrine neoplasia type 1 (MEN1) is a rare tumor syndrome caused by mutations in the *MEN1* gene. It has an autosomal dominant inheritance pattern that is approximately 95% penetrant at 40-50 years and shows no gender bias Its main clinical features are multi-gland parathyroid adenomas (90-100% of patients), anterior pituitary adenomas (30-40%), and gastrinomas/PanNETs (30-40%)^4,5^. A retrospective study of 576 germline *MEN1* mutations in patients with MEN1 syndrome showed variants scattered throughout the coding region. In this study, 69% of pathologic mutations were truncations^6^. Mutations in the *MEN1* gene are found in sporadic tumors as well. Sequencing of tumors from prostate cancer patients within intermediate- and high-risk groups showed mutations in the *MEN1* gene (1/8 patients, T3bNXM0)^7^. Additionally, lung carcinoid tumors contain *MEN1* mutations in approximately 17% of cases^8-10^. In one study, decreased *MEN1* gene expression correlated with the presence of mutations, occurrence of distant metastasis, and decreased overall survival^8^.

The majority of PanNETs develop sporadically and recent genomic analysis of primary tumors revealed a complex genomic landscape with four common oncogenic events. The tumor suppressor gene *MEN1* was found to be mutated in approximately 44% of sporadic PanNETs^11^. The *MEN1* gene is comprised of 10 exons and its protein, Menin, is translated from exons 2-10 into a 610 amino acid protein. Two nuclear localization signals are found on the C-terminal of Menin that are important for its localization in the nucleus^12,13^. Menin functions as a member of COMPASS-like complexes containing MLL and MLL2 that are responsible for methylation of H3K4 and regulation of approximately 5% of global H3K4 trimethylation (H3K4me3) levels^14^. Its role within the complex is required for localization of the complex to chromatin at specific gene promoters^15,16^; however, the signaling pathways affected by *MEN1* mutations in PanNETs remains unknown. Therefore, we initiated a pan-genomic analysis of primary pancreatic neuroendocrine tumors to determine the association between mutations and signaling pathways and elucidate which are responsible for PanNET initiation and progression. We identified several signaling pathways that are deregulated downstream of mutant *MEN1* in a set of sporadic primary pancreatic neuroendocrine tumors.

## Materials and Methods Patients and tissue samples

A set of formalin-fixed, paraffin-embedded sporadic pancreatic neuroendocrine tumors from de-identified patients who underwent surgical resection at Banner University Medical Center (2005-2019) without receiving preoperative therapy was utilized for this study. Hematoxylin and eosin stained slides were reviewed on all cases along with corresponding surgical pathology reports. Functional status, stage, grade, and Ki-67 proliferation index was determined based on the surgical pathology report. This study was approved by the Institutional Review Board of The University of Arizona.

### Targeted exome sequencing

Genomic DNA was isolated from FFPE tissue sections containing greater than 85% tumor cellularity using the Qiagen QIA Amp FFPE Tissue kit (Qiagen, Valencia, CA). 40 ng of DNA from each sample was used for deep sequencing using the Ion Ampliseq Comprehensive Cancer Panel (Thermo Fisher Scientific, Waltham, MA). Barcoded libraries (16,000 amplicons distributed among four pools) were generated from 40 ng of DNA using the Ion Ampliseq library 2.0 kit. Quality of libraries was determined using an Agilent 2100 bioanalyzer instrument (Agilent Technologies, Santa Clara, CA) and quantity was measured by qPCR using the Ion Library Quantification kit. Amplified and clonal templates from libraries were generated by emulsion PCR using the Ion Proton Template OT2 kit. Sequencing was performed by multiplexing four barcoded templates on each Proton I chip on the Ion Torrent instrument. All experiments were performed according to the manufacturer’s instructions. Unaligned binary data files generated by the Ion Torrent instrument were uploaded into Ion Reporter and analyzed using default settings. Alignment of the sequencing reads to the reference genome (hg19) was performed using the Ion Torrent Suite software. Identification of sequence variants was performed by the Ion Torrent Variant Caller software and coverage of each amplicon was determined using the Coverage Analysis software. Integrative Genomics Viewer^17^ was used to visualize the read alignment and presence of variants against the reference genome as well as to confirm variant calls by checking for strand bias and sequencing errors. Variants were filtered based on the following criteria: non-synonymous, exonic, p-value ≤ 0.05, greater than 15X coverage, and an alternate allele frequency ≥ 10%.

### Microarray

Total RNA was isolated from FFPE tissue sections containing greater than 85% tumor cellularity using the Qiagen miRNeasy FFPE kit (Qiagen, Valencia, CA) and quantified by absorption measurements at 260 nm. 10ng of RNA from each sample was used for gene expression analysis using the Affymetrix Clariom D arrays (Thermo Fisher Scientific, Waltham, MA). Arrays were read using the Affymetrix 2500A scanner (Affymetrix, Santa Clara, CA) and analyzed using the Affymetrix Transcriptome Analysis Console version 4.0 (Thermo Fisher Scientific, Waltham, MA). All experiments were performed according to the manufacturer’s instructions. Raw data (CEL files) were summarized to both the gene- and transcript-level signal and normalized using the SST-RMA algorithm to reduce background and increase fold change. Samples were labeled according to their *MEN1* or *FGFR3* mutation status (wild-type or mutant) and each transcript was normalized to its median. Exploratory grouping analysis was performed to determine clustering of samples, batch effects determined variation between samples, and limma^18^ was performed for differential expression analysis. Genes that were considered significant (p-value ≤ 0.05) and had a fold change either ≥ 5 or ≤ −5 were considered for further analysis.

### Quantitative reverse-transcription PCR

Total RNA was isolated from FFPE tissue sections containing greater than 85% tumor cellularity using the Qiagen miRNeasy FFPE kit (Qiagen, Valencia, CA) and quantified by absorption measurements at 260 nm. 1 μg of RNA was reverse transcribed using 1 μg/ml random hexamer primers and Superscript II reverse transcriptase (Thermo Fisher Scientific, Waltham, MA). Gene expression was performed using TaqMan primer/probe sets containing a FAM quencher to *MEN1* (Hs00365720_m1), *ITGA2* (Hs00158127_m1), *EZH2* (Hs00544830_m1), *CDH1* (Hs01023895_m1), *ALK* (Hs01058318_m1), *VEGFA* (Hs00900055_m1), and *HPRT* (Hs02800695_m1) (Thermo Fisher Scientific, Waltham, MA). qPCR was performed using the TaqMan Universal Master Mix, no UNG on an ABI Prism 7500 Sequence Detection System (Thermo Fisher Scientific, Waltham, MA). Differences in expression between *MEN1* wild-type and *MEN1* mutant tumors was determined using the comparative Ct method described in the ABI user manual relative to *HPRT*. Differences in expression between *FGFR3* wild-type tumors and *FGFR3* mutant tumors was determined using the comparative Ct method relative to *HPRT*. Total RNA was isolated from each tumor in duplicate and each RNA sample was processed by PCR in triplicate.

### Immunohistochemistry

Tumor tissue sections were deparaffinized with Xylene and rehydrated. Antigen retrieval was carried out with antigen unmasking solution (Vector Laboratories, Burlingame, CA) at 95-100°C for 30 minutes. Slides were incubated in blocking solution (3% normal donkey serum, 1% Tween20 in PBS) for 1 hour at room temperature. Primary antibody in blocking solution was added and slides were incubated overnight at 4°C. The primary antibodies for SYP (ab8049, 1:1000), Menin (ab2605, 1:1000), ITGA2 (ab133557, 1:1000) and EZH2 (ab191080, 1:1000) were purchased from Abcam (Cambridge, United Kingdom). The primary antibody to H3K27me3 (#07-449, 1:1000) was purchased from Millipore (Burlington, MA). The primary antibody to total FGFR3 (sc-13121, clone B-9, 1:1000) was purchased from Santa Cruz Biotechnology (Dallas, TX). The primary antibodies to E-cadherin (#14472, clone 4A2, 1:500) and VE-cadherin (#2158, 1:250) were purchased from Cell Signaling Technology (Danvers, MA). Biotinylated secondary antibody was added and incubated for 30 minutes followed by incubation with Vectastain Elite avidin biotin complex for an additional 30 minutes (Vector Laboratories, Burlingame, CA). Slides were developed using the SignalStain DAB Substrate kit (Cell Signaling Technology, Danvers, MA) and counterstained with hematoxylin. The immunohistochemical stained slides were scanned at 10X and 40X. To determine the difference in protein localization in *MEN1*/*FGFR3* mutant tumors, membrane, cytoplasmic, and nuclear staining was counted and displayed as percent stained divided by the total number of cells. Differences in the percentage of membrane, cytoplasmic, and nuclear staining was analyzed by a student’s t-test. A p-value ≤ 0.05 was considered statistically significant.

### Analysis of PanNET variants

Variants selected from targeted exome sequencing were analyzed using the software MUtations For Functional Impact on Network Neighbors (MUFFINN)^19^. MUFFINN is a publicly available method (https://www.inetbio.org/muffinn/) that utilizes a protein-protein interaction network to identify neighborhoods of genes that are affected by a user’s input list of variants. For this project, we utilized the direct neighbor max (DNmax) algorithm which measures the impact of the most frequently mutated neighbor as opposed to the sum of all directly mutated neighbors. The Search Tool for the Retrieval of Interacting Genes/Proteins (STRING) database of known protein-protein interactions was utilized to develop gene networks in both *MEN1* or *FGFR3* wild-type PanNETs and *MEN1* or *FGFR3* mutant PanNETs using the gene lists outputted by MUFFINN.

## Results

To better understand the biological landscape of PanNETs, we performed targeted exome sequencing on a panel of oncogenes and tumor suppressor genes known to be involved in multiple cancers using the Ion Torrent Comprehensive Cancer Panel. We randomly chose six well-characterized sporadic PanNETs from patients who had not received therapy prior to surgical resection (Table 1). In order to maintain high neoplastic cellularity, we stained our tumor samples with Synaptophysin (SYP) to ensure enrichment of neuroendocrine cells for targeted exome sequencing (data not shown). Genomic DNA from the six samples was used to prepare fragment libraries suitable for sequencing. Coding sequences were enriched by emulsion PCR using the Ion Proton Template OT2 kit. Sequencing was performed by multiplexing four barcoded templates on each Proton I chip on the Ion Torrent instrument. The average coverage of each base of the targeted regions was 1190-fold with an average rate of on-target reads equal to 84%.

**Table 1.** Clinical and pathological characteristics of the six pancreatic neuroendocrine tumors randomly selected for targeted exome sequencing.

|  | PanNET Samples (N = 6) |
| --- | --- |
| Age - Median (Range) | 69.5 (38 – 84) |
| Gender – Number (%) |  |
| Male | 3 (50%) |
| Female | 3 (50%) |
| **Functional Status – Number (%)** |  |
| Functional | 1 (17%) |
| Non-Functional | 5 (83%) |
| **Tumor Site – Number (%)** |  |
| Head | 3 (50%) |
| Body | 3 (50%) |
| Tail | 1 (17%) |
| **TNM Staging – Number (%)** |  |
| Stage 1 | 0 (0%) |
| Stage 2 | 2 (33%) |
| Stage 3 | 4 (66%) |
| Stage 4 | 0 (0%) |
| **KI67 Labeling Index – Number (%)** |  |
| < 2% | 1 (17%) |
| ≤ 2-20% | 4 (66%) |
| > 20% | 1 (17%) |
| **Metastatic Site – Number (%)** |  |
| Lymph Node | 3 (50%) |
| Liver | 2 (33%) |
| Other | 1 (17%) |
| None | 3 (50%) |
| **Co-Morbidity – Number (%)** |  |
| Chronic Pancreatitis | 4 (66%) |
| Perineural Invasion | 2 (33%) |
| Chronic Cholecystitis | 2 (33%) |
| Other | 3 (50%) |
| None | 0 (0%) |

Overall, we identified 103 mutations in 76 genes amongst the six PanNET tumors. The mutations per tumor ranged from 8 to 31 with a mean of 17. Sequencing revealed mutations in genes that have been previously found in other genomic PanNET studies^11,20^: 3/6 contained a mutation in the *MEN1* gene, 2/6 showed mutations in the *MTOR* and *TSC2* genes, and 1/6 had a mutation in *ATRX* (Figure 1a). Subsequently, we examined other genes mutated in our primary PanNETs. Similar to pulmonary carcinoids^9^, our PanNETs contained mutations in various chromatin modifying genes including *EZH2*, histone lysine demethylases, histone lysine methyltransferases, histone acetylation, and chromatin remodeling genes including *ARID1A* (Figure 1a). Interestingly, 5/6 tumors contained a mutation in a receptor tyrosine kinase and 3/6 showed mutations in *PARP1* (Figure 1a). A list of specific point mutations can be found in Table 2. These data suggest a potential cooperation between alterations in epigenetic modulators along with conventional genetic alterations being sufficient to drive initiation and progression of pancreatic neuroendocrine tumors.

**Table 2.** Point mutations identified by targeted exome sequencing within our set of randomly selected pancreatic neuroendocrine tumors.

| Sample Name | Age | Sex | Stage | KI67 | Histone Covalent Modifiers | SWI/SNF Complex | Recurrently Mutated Genes in PanNETs | Other Interesting Genes |
| --- | --- | --- | --- | --- | --- | --- | --- | --- |
| R150231 | 84 | M | T2N0MX | < 2% | EP300_p.His2324fs<br>EP300_p.Pro2329fs<br>KDM6A_p.Pro1395Ala |  |  | AXL_p.Pro833Arg |
| R150280 | 70 | M | T3N1M1 | 60% | CREBBP_p.Arg1347Pro<br>KMT2D_p.His313Pro | ARID1A_p.Pro418Thr |  | ALK_p.Trp593Cys<br>MAPK1_p.Tyr193Asn<br>PARP1_p.Gly135Asp |
| R160068 | 69 | F | T2N1M1 | 7% | EP300_p.Gly269Val<br>EP300_p.Ala2356Asp<br>EZH2_p.Ala581Ser<br>KMT2D_p.Gln3720Leu<br>PSIP1_p.Asp234His |  | TSC2_p.Ala623Ser<br>TSC2_p.Phe1275Val | MSH6_p.Ser227Arg<br>PARP1_p.Gln353fs<br>SRC_p.Thr304Arg |
| R160018 | 53 | F | T3N1MX | 15% | KAT6B_p.Gly2011fs<br>MEN1_p.Ala100_Gly110del |  | MTOR_p.Thr600Ile | DICER1_p.Arg490Cys<br>FGFR3_p.Pro587Thr |
| G120029 | 38 | F | T2N0M1 | 5% | KAT6B_p.Ser124Phe<br>MEN1_p.Arg237Cys |  |  | FGFR3_p.Val682Ile |
| G130043 | 83 | M | T3N0MX | ≤ 2% | MEN1_p.Trp346Ter |  | ATRX_p.Val2180Leu | PARP1_p.Gln353fs |

**Table 3.** Percentage of membrane, cytoplasmic, and nuclear staining in adjacent normal, *MEN1* wild-type, and *MEN1* mutant tumors for ITGA2, EZH2, and E-cadherin. To determine the difference in protein localization in *MEN1* mutant tumors, membrane, cytoplasmic, and nuclear staining was counted and displayed as a percentage (number of cells stained divided by the total number of cells).

|  | % membrane | % cytoplasmic | % nuclear |
| --- | --- | --- | --- |
| **Adjacent Normal** |  |  |  |
| ITGA2 | 0.86 | 10.42 | 20.85 |
| EZH2 | 0.00 | 10.91 | 17.36 |
| E-cadherin | 95.66 | 4.50 | 1.04 |
| ***MEN1* wild-type** |  |  |  |
| ITGA2 | 0.50 | 58.04 | 22.81 |
| EZH2 | 0.00 | 25.26 | 38.36 |
| E-cadherin | 41.24 | 75.02 | 49.96 |
| ***MEN1* mutant** |  |  |  |
| ITGA2 | 0.45 | 93.82 | 88.87 |
| EZH2 | 0.00 | 68.57 | 72.10 |
| E-cadherin | 0.00 | 71.72 | 43.87 |

**Figure 1.**
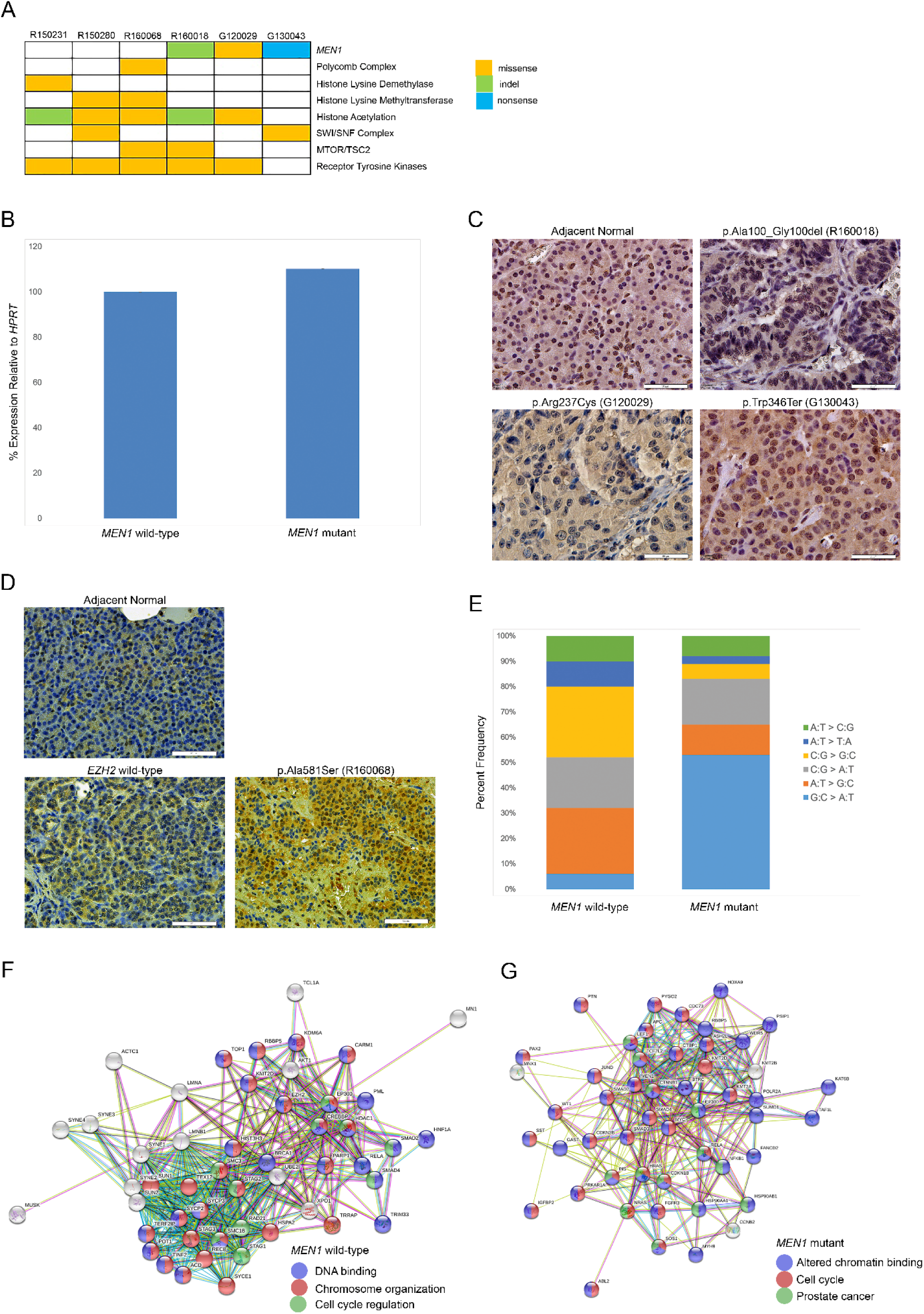
Targeted exome sequencing reveals mutational variants in pancreatic neuroendocrine tumors and novel signaling pathways deregulated downstream of mutant *MEN1*. (a) Mutational analysis of PanNETs. Targeted exome sequencing using the Ion Torrent Comprehensive Cancer Panel identified mutational variants within several genes/pathways. Patients are denoted in each individual column while genes/pathways are indicated in each row. Missense mutations are colored yellow, insertions and deletions are depicted in green, and nonsense mutations are shown in blue. (b) Quantitative PCR for *MEN1* expression. For analysis, Ct values were averaged across *MEN1* wild-type tumors and *MEN1* mutant tumors. (c) Immunohistochemistry for Menin protein. FFPE tissue sections were stained with anti-Menin overnight and visualized with DAB. Images were acquired at 60X and the white bar denotes a scale of 50 μm. (d) Immunohistochemistry for H3K27me3 protein. FFPE tissue sections were stained with anti-H3K27me3 overnight and visualized with DAB. Images were acquired at 40X and the white bar denotes a scale of 50 μm. (e) COSMIC mutational signatures of PanNETs. To determine the difference in single base substitution subtypes, each subtype was counted and displayed as a percentage (number of single base substitutions for each subtype divided by the total number of single base substitutions). (f) MUFFINN analysis of *MEN1* wild-type PanNETs. Genes associated with the top three biological processes associated with these tumors are shown in blue, red, and green. (g) MUFFINN analysis of *MEN1* mutant PanNETs. Genes associated with the top three biological processes associated with these tumors are shown in blue, red, and green.

The *MEN1* mutations found in our PanNETs were p.Ala100_Gly110del (R160018), p.Arg237Cys (G120029), and p.Trp346Ter (G130043). Cross-checking our *MEN1* variants with UniProt and the UMD-MEN1 mutation database (http://www.umd.be/MEN1/), we determined that our variants do not occur naturally and have yet to be described in cancer; however, two of the mutations (p.Arg237Cys and p.Trp346Ter) occur within a region that has been shown to interact with FANCD2^21^. Quantitative PCR of our *MEN1* wild-type and *MEN1* mutant PanNETs showed no difference in *MEN1* RNA expression (Figure 1b). To determine if our novel *MEN1* mutations were functional, we performed immunohistochemistry (IHC) using a Menin antibody to determine changes in Menin expression and localization in our *MEN1* mutant PanNETs (Figure 1c). Menin stained strongly positive in approximately 50% of nuclei in adjacent normal pancreas tissue with no cytoplasmic staining seen. R160018 (p.Ala100_Gly110del) stained similar to the adjacent normal tissue. G120029 (p.Arg237Cys) showed weak Menin staining in both the nuclear and cytoplasmic compartments while G130043 (p.Trp346Ter) stained strongly positive in approximately 50% of nuclei and showed moderate positive staining in the cytoplasm in the majority of cells. These data suggest that the p.Arg237Cys and p.Trp346Ter mutations may be functional by altering Menin expression and protein localization, respectively.

The *EZH2* mutation found in one of our PanNETs was Ala581Ser (R160068) and has not been previously described. Our *EZH2* variant occurs within the conserved CXC domain located immediately N-terminal to the SET domain. The CXC domain is unique to *EZH2* and is coordinated by two clusters of three zinc ions. Mutations within the first and second zinc ions of the CXC domain (His525Asn and Cys571Asn, respectively) have been reported in acute myeloid leukemia and myelodysplastic syndrome suggesting a role for disruption of the *EZH2* CXC domain in cancer^22,23^. To determine EZH2 activity as a result of this mutation, IHC using an antibody to H3K27me3 was performed (Figure 1d). In adjacent normal pancreas, approximately one-third of nuclei in the acinar compartment showed strong positive staining while the *EZH2* wild-type tumor showed weak nuclear staining in less than 25% of nuclei. Conversely, the *EZH2* mutant tumor showed strong positive staining in approximately 50% of nuclei suggesting that this mutation may be functional in *EZH2* activation.

Next, we separated our PanNET tumors based on their *MEN1* status (wild-type or mutant) to examine mutational differences between the two groups. The *MEN1* wild-type tumors contained 61 mutations in 51 genes (range = 11 to 31, mean = 20) while the *MEN1* mutant tumors showed 42 mutations in 31 genes (range = 8 to 24, mean = 14). There were no dissimilarities in the type of mutations based on the *MEN1* mutation status in PanNETs (data not shown). Interestingly, there were differences in the mutational signature between the two groups. The *MEN1* mutant tumors showed a higher proportion of G:C > A:T and C:G > A:T mutations compared to wild-type tumors. (Figure 1e). This mutational signature is reminiscent of COSMIC mutational signature 6 which has been associated with DNA mismatch repair deficiency^24^. These data suggest that *MEN1* mutant PanNETs may arise through a separate mechanism than their wild-type counterparts.

A major challenge of exome sequencing is to identify the significance of infrequent mutations, particularly in smaller patient populations. Therefore, we utilized MUFFINN to identify neighborhoods of genes that are impacted by mutations in *MEN1*. Using the top 50 genes generated by MUFFINN for each group (*MEN1* wild-type vs. *MEN1* mutant), we found the signaling pathways regulated by both wild-type and mutant *MEN1* in our PanNETs. Not surprisingly, mutations in *MEN1* wild-type tumors regulated known biological processes associated with Menin including: chromosome organization (red), DNA binding (blue), and cell cycle regulation (green) (Figure 1f). Mutations in *MEN1* mutant PanNETs, however, showed deregulation in cell cycle (red), altered protein binding (blue), and signaling pathways associated with prostate cancer (green) including members of the PI3K/AKT pathway (Figure 1g). This suggests that mutations in the *MEN1* gene lead to increased cell proliferation potentially through the PI3K/AKT pathway.

Although the genomic landscape of pancreatic neuroendocrine tumors has been established by several groups^11,25,26^, the transcriptomic landscape of this disease remains largely unknown. Therefore, we analyzed our primary PanNET samples via microarray to determine which genes are upregulated or downregulated based on the tumor’s *MEN1* status to gain insight into the pathways deregulated downstream of mutant *MEN1*. When compared to wild-type *MEN1* tumors, mutant *MEN1* PanNETs showed deregulation of 229 genes (63% upregulated and 37% downregulated). Hierarchical clustering confirmed that *MEN1* wild-type and *MEN1* mutants PanNETs have distinct gene expression patterns (Figure 2a). Notable upregulated genes in *MEN1* mutant tumors include *ITGA2* (7-fold) and *EZH2* (11-fold) (Figure 2b). Additionally, *CDH1* (E-cadherin) was found to be downregulated in mutant *MEN1* PanNETs (−18-fold) (Figure 2b). Interestingly, upon pathway analysis, *MEN1* mutant PanNETs showed potential deregulation in genes involved in epithelial-to-mesenchymal transition (upregulated: *LRP6, AKT2*, and *EZH2*; downregulated: *CDH1*)^27-32^. To validate our microarray results, we used quantitative PCR (qPCR) using TaqMan probes to *ITGA2, EZH2*, and E-cadherin which have been shown to be involved in cancer progression^33,34^ and cell invasion^35,36^. When compared with wild-type PanNETs, tumors containing mutant *MEN1* showed a 6-fold increase in *ITGA2* expression, a 2-fold increase in *EZH2*, and a 75% decrease in *CDH1* expression (Figure 2c). This suggests that PanNETs containing *MEN1* mutations may be more aggressive and invasive than their wild-type counterparts.

**Figure 2.**
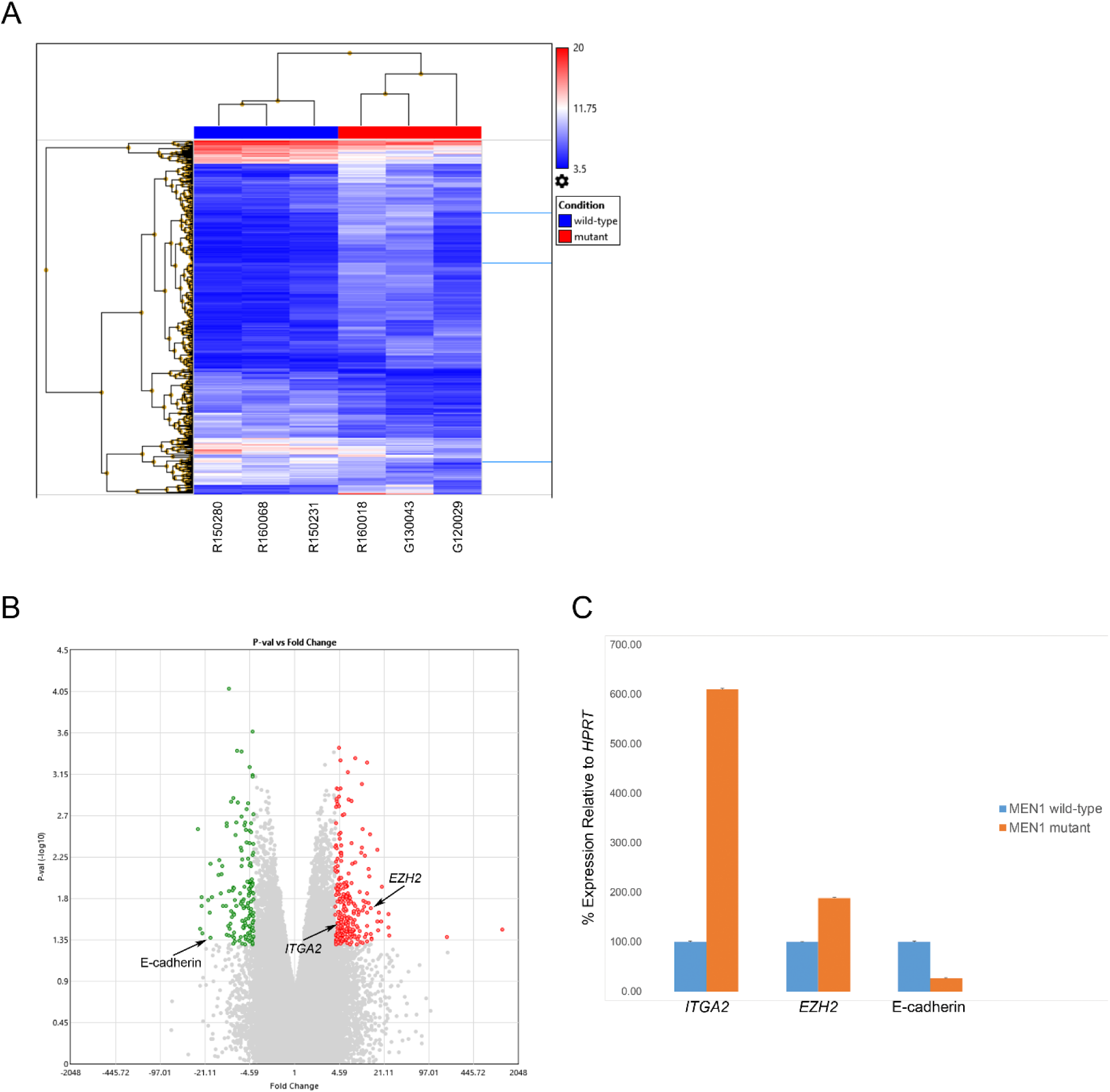
Transcriptome analysis in *MEN1* mutant pancreatic neuroendocrine tumors reveals deregulated expression in genes involved in epithelial-to-mesenchymal transition. (a) Hierarchical clustering of *MEN1* mutant PanNETs vs. *MEN1* wild-type PanNETs. (b) Volcano plot of *MEN1* mutant PanNETs vs. *MEN1* wild-type PanNETs. Each dot represents one gene. Green dots denote genes that are downregulated in *MEN1* mutant tumors while red dots indicate upregulated genes in *MEN1* mutant tumors. Genes chosen for further validation are indicated with arrows. (c) Quantitative PCR of selected genes.

Immunohistochemistry for ITGA2 showed a statistically significant increase in nuclear staining in *MEN1* mutant PanNETs compared to *MEN1* wild-type PanNETs (p = 0.0047) and adjacent normal (p = 0.001) (Figure 3a and 4). IHC for E-cadherin showed a statistically significant decrease in membrane E-cadherin localization in *MEN1* mutant PanNETs compared to *MEN1* wild-type PanNETs (p = 0.0292) and adjacent normal (p = 5.2827×10^−5^) (Figure 3b and 4). IHC for EZH2 showed a statistically significant increase in cytoplasmic localization in *MEN1* mutant PanNETs compared to *MEN1* wild-type PanNETs (p = 0.0071) and adjacent normal (p = 0.0001) (Figure 3c and 4). Additionally, a statistically significant increase in nuclear EZH2 localization in *MEN1* mutant PanNETs was seen as well (Figure 3c and 4). No differences in staining intensity was seen for ITGA2, EZH2, or E-cadherin. These results further validate our microarray data and suggest that PanNETs containing *MEN1* mutations may be more aggressive and invasive than their wild-type counterparts.

**Figure 3.**
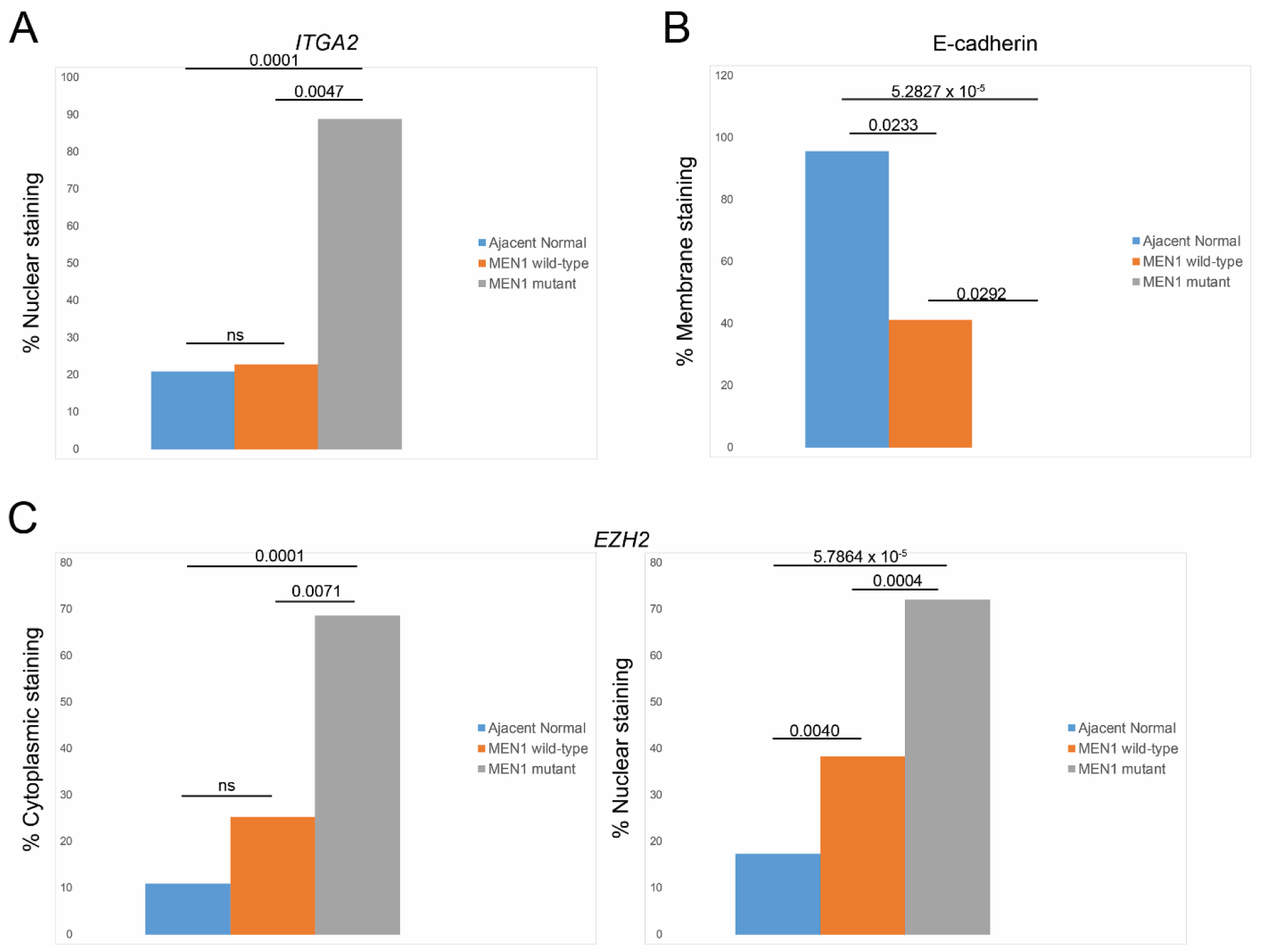
Changes in protein localization between *MEN1* wild-type PanNETs and *MEN1* mutant PanNETs. To determine the difference in protein localization in *MEN1* mutant tumors, membrane, cytoplasmic, and nuclear staining was counted and displayed as a percentage (number of cells stained divided by the total number of cells). Differences in the percentage of membrane, cytoplasmic, and nuclear staining was analyzed by a student’s t-test. A p-value ≤ 0.05 was considered statistically significant.

Two-thirds of our mutant *MEN1* PanNETs also contained a mutation in *FGFR3*, an evolutionarily conserved receptor tyrosine kinase. Upon ligand binding, FGFR3 dimerizes leading to autophosphorylation of the cytoplasmic kinase domain. The two main substrates of *FGFR3* are PLCγ and FRS2. Phosphorylation of FRS2 is essential for activation of the MAPK pathway downstream of the receptor^37,38^ while a separate complex involving GAB1 leads to the recruitment of PI3K leading to activation of the AKT anti-apoptotic pathway^39^. Germline mutations in the *FGFR3* kinase domain (Ile538, Asn540, and Lys650) have been shown to increase catalytic activity independently of receptor dimerization in three separate dwarfing syndromes with varying clinical severity^40,41^. Activating mutation Lys650Glu in *FGFR3* has been shown in multiple myeloma^42^ and overexpression of *FGFR3* occurs in approximately 92% of bladder cancer and 93% of cervical carcinomas^43^.

Two out of our six samples contained mutations in the FGFR3 kinase domain. Sample R160018 had a Pro587Thr mutation while sample G120029 had a Val682Ile mutation. According to Uniprot, neither of these *FGFR3* somatic mutations have been described as natural variants suggesting that they may play a role in PanNET progression. To determine if our novel *FGFR3* mutations were functional, we performed IHC using a total FGFR3 antibody to determine changes in total FGFR3 expression and localization in our *FGFR3* mutant PanNETs (Figure 5a). FGFR3 stained strongly positive in the cytoplasm of the Islet of Langerhans, was weakly positive in the cytoplasm of approximately 50% of the acinar cell compartment, and was moderately positive in the nucleus of approximately 10% of the acinar cell compartment in adjacent normal pancreas tissue. G120029 (p.Val682Ile) showed moderate cytoplasmic staining in all cells and weak nuclear staining in a small number of cells while R160018 (p.Pro587Thr) showed moderate cytoplasmic staining in all cells, strong nuclear staining in approximately 10% of cells, and moderate nuclear staining in 50% of cells. These data suggest that the Val682Ile mutation may not be functional while the Pro587Thr mutation may activate the receptor.

**Figure 4.**
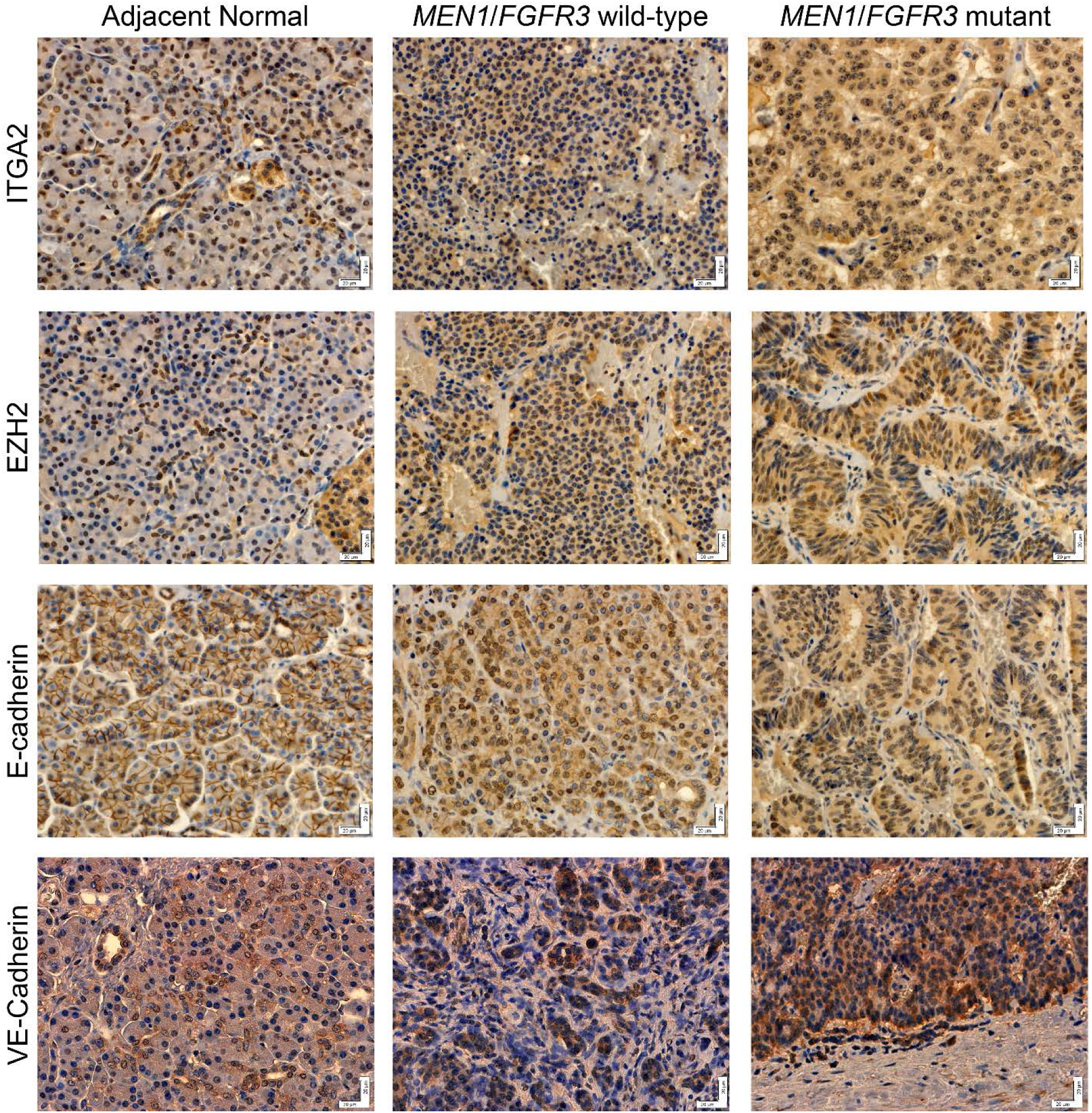
Deregulated protein expression and localization in *MEN1*/*FGFR3* mutant PanNETs. Immunohistochemistry for ITGA2, EZH2, E-cadherin, and VE-Cadherin proteins. FFPE tissue sections were stained with primary antibody overnight and visualized with DAB. Images were acquired at 40X and the white bar denotes a scale of 20 μm.

**Figure 5.**
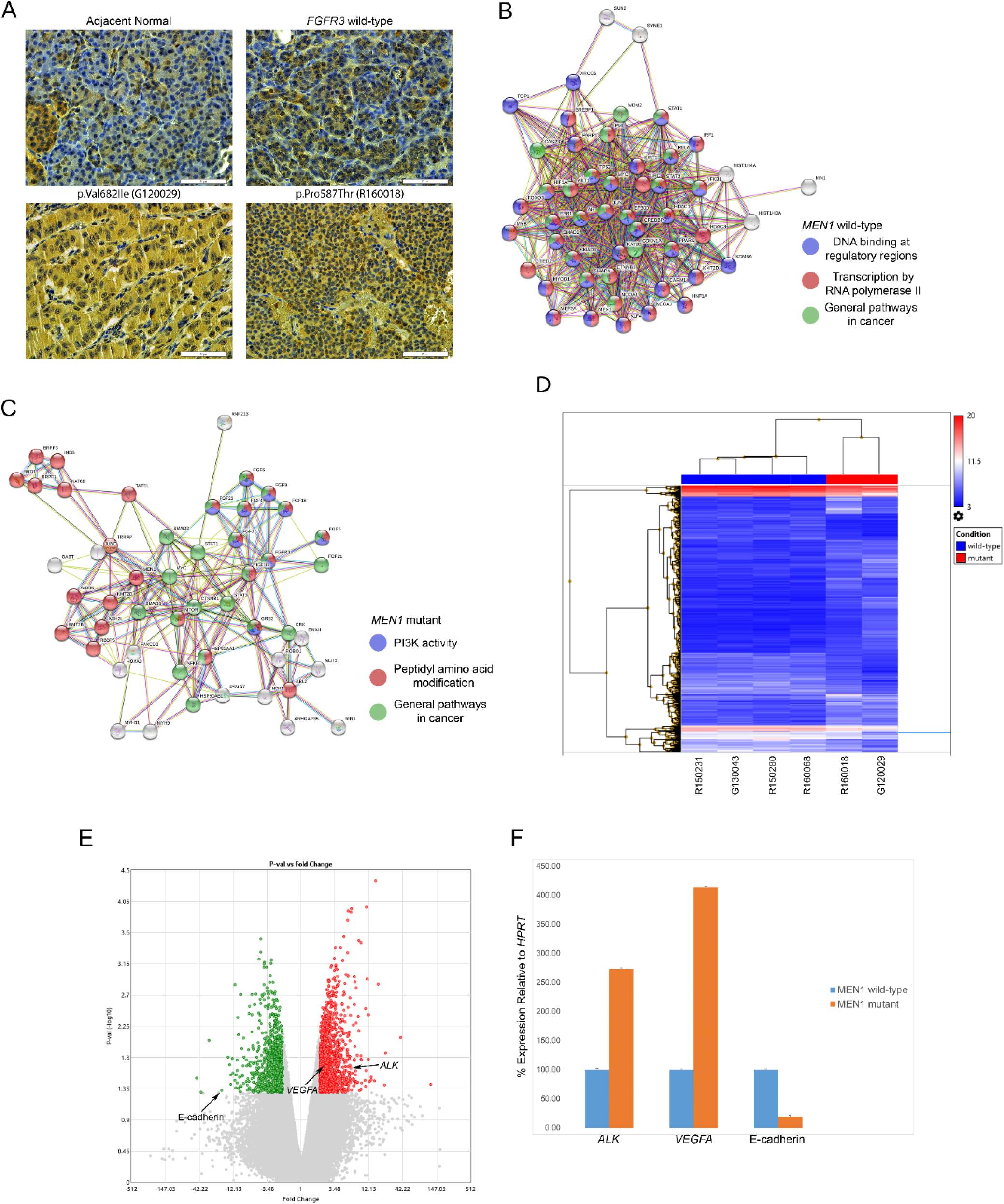
Transcriptome analysis in *FGFR3* mutant pancreatic neuroendocrine tumors reveals deregulated expression in genes involved in angiogenesis. (a) Immunohistochemistry for total FGFR3 protein. FFPE tissue sections were stained with anti-FGFR3 overnight and visualized with DAB. Images were acquired at 40X and the white bar denotes a scale of 50 μm. (b) MUFFINN analysis of *FGFR3* wild-type PanNETs. Genes associated with the top three biological processes associated with these tumors are shown in blue, red, and green. (c) MUFFINN analysis of *FGFR3* mutant PanNETs. Genes associated with the top three biological processes associated with these tumors are shown in blue, red, and green. (d) Hierarchical clustering of *FGFR3* mutant PanNETs vs. *FGFR3* wild-type PanNETs. (e) Volcano plot of *FGFR3* mutant PanNETs vs. *FGFR3* wild-type PanNETs. Each dot represents one gene. Green dots denote genes that are downregulated in *FGFR3* mutant tumors while red dots indicate upregulated genes in *FGFR3* mutant tumors. Genes chosen for further validation are indicated with arrows. (f) Quantitative PCR of selected genes.

Separating our PanNETs based on their *FGFR3* mutational status, we applied MUFFINN to determine which signaling pathways are regulated by *FGFR3* in our wild-type and mutant tumors. Mutations in *FGFR3* wild-type tumors were shown to regulate transcription by RNA polymerase II (red), DNA binding at regulatory regions (blue), and general pathways involved in cancer (green) (Figure 5b). Conversely, mutations in *FGFR3* mutant PanNETs showed deregulation in peptidyl-amino acid modification (red), PI3K activity (blue), and general pathways involved in cancer (green) (Figure 5c). These results suggest that *FGFR3* mutations may promote PanNET progression through deregulated PI3K signaling.

Next, we looked at our expression data to determine pathways deregulated downstream of *FGFR3*. When compared to wild-type tumors, *FGFR3* mutant PanNETs showed downregulation of 200 genes (55% upregulated and 45% downregulated). Hierarchical clustering confirmed that *FGFR3* wild-type and *FGFR3* mutant PanNETs have distinct gene expression patterns (Figure 5d). Notable upregulated genes in *FGFR3* mutant tumors include *VEGFA* (7-fold) and *ALK* (10-fold) (Figure 5e). Additionally, *CDH1* was found to be downregulated in mutant *FGFR3* PanNETs (−12-fold) (Figure 5e). To validate our microarray results, we performed qPCR using TaqMan probes to *VEGFA, ALK*, and *CDH1* which have been shown to be involved in angiogenesis^44-46^. When compared with wild-type PanNETs, tumors containing mutant *FGFR3* showed a 2.7-fold increase in *ALK* expression, a 4-fold increase in *VEGFA* expression, and an 80% decrease in *CDH1* expression (Figure 5f). Immunohistochemistry for VE-Cadherin showed an increase in the number of blood vessels present in both *FGFR3* wild-type and *FGFR3* mutant tumors; however, due to differences in tissue sectioning, variances in the number of blood vessels between *FGFR3* wild-type and *FGFR3* mutant tumors could not be drawn (Figure 4). These results suggest that PanNETs containing *FGFR3* mutations may lead to more angiogenic tumors.

## Discussion

Several genomic studies looking at the genetic drivers of PanNETs have shown that PanNETs do not harbor traditional driver mutations in oncogenes and tumor suppressor genes that have been identified in other cancer types^11,25,26^. Instead, PanNETs show genetic variation in multiple classes of epigenetic modifying genes, including *MEN1*. While epigenetics has been shown to play a major role in cancer development^47^, it still remains only “one hit” of Knudson’s “two-hit” hypothesis for cancer formation. Our sequencing results show that in addition to mutations in *MEN1* and other epigenetic regulators, mutations in genes shown to be involved in the initiation of other cancers were present as well. Therefore, our results suggest a potential cooperation between alterations in epigenetic regulators, including *MEN1*, and other conventional alterations as being sufficient to drive the transformation of pancreatic neuroendocrine tumors.

Somatic mutations are present in every cell within the human body and accumulate throughout our lifetime. They are the consequence of multiple different processes from deficiencies in DNA repair to enzymatic modification of DNA to exogenous mutagens. Each of these different mutational processes generate a unique combination of mutations termed “mutational signatures”. Therefore, these signatures can be used to probe underlying causes of cancer formation in tumors. We found that our *MEN1* mutant tumors contained more G:C > A:T and C:G > A:T mutations than their wild-type counterparts. This mutational signature is similar to the COSMIC Mutational Signature 6 which has been shown to be enriched in tumors associated with DNA mismatch repair deficiencies. While mutations in mismatch repair genes were not present in any of our *MEN1* mutant PanNET tumors, we cannot rule out potential epigenetic silencing of mismatch repair genes which has been shown to be involved in multiple other tumor types^48-53^.

Normal epithelial cells regulate growth through tightly regulated systems using specific proteins as checks and balances to ensure cell proliferation occurs only when necessary. Cancer cells have evolved to exploit these systems in a variety of ways including increased growth and proliferation signals along with the inhibition of cell cycle checkpoints. FGFR3 signals through a complex involving GAB1 leading to the recruitment of PI3K and activation of the AKT pro-proliferation pathway^39^. On the other hand, Menin directs the COMPASS-like complex to the promoters of p18 and p27 to inhibit cell cycle progression^54-56^. Our sequencing results show that *MEN1* and *FGFR3* mutations occur together and show mutational enrichment of pathways involving members of the PI3K/AKT signaling pathway. While our string analysis does not show any direct interaction between *MEN1* and *FGFR3*, it is possible that the two mutations act in tandem to promote cell proliferation. Initiation of the “go” signal may come from constitutively active FGFR3 while loss of the “stop” signal may come from the absence of Menin at the p18 and p27 promoters.

The FGFR family of receptor tyrosine kinases share a high percentage of sequence homology (50-72%) to one another^57^. One of our *FGFR3* mutant PanNETs (G120029) contained the mutation Val682Ile. While this mutation is situated in the kinase domain of the FGFR3 protein, protein alignment between FGFR3 and FGFR4 shows that Ile is the amino acid at this position in wild-type FGFR4^58^ suggesting that this mutation is most likely benign and does not contribute to this tumor. Interestingly, the majority of receptor tyrosine kinases split the canonical kinase domain with a non-homologous sequence of amino acids termed the kinase insert domain^59^. FGFRs contain a “small” kinase insert domain of 37 amino acids. Upon protein crystallization, this kinase insert domain forms a disordered loop that is hard to crystalize^58,60^. It has been shown that phosphorylation of Tyr583 and Tyr585 in the kinase insert domain of FGFR1 is necessary for activation of mitogenesis through the receptor^61^. Additionally, mass spec showed heterogeneous phosphorylation of Tyr586 in the kinase insert domain of FGFR2^58^; however, the authors did not describe any functionality associated with this post-translational modification. The second *FGFR3* mutation found in our PanNETs (R160018) was Pro587Thr which occurs within this disordered loop. While IHC of this tumor showed more nuclear localization of FGFR3 which has been associated with poor overall survival in pancreatic ductal adenocarcinoma^62^, it remains to be determined if this mutation has any functional impact on the biology of this tumor.

Cell differentiation is the process in which progenitor cells mature into cells with specific roles. In cancer, cells become de-differentiated and start to lose their normal architecture and function. Well-differentiated cancer cells tend to look more like the normal tissue and grow more slowly and are less invasive than poorly differentiated or undifferentiated cells. EZH2 has long been known to induce de-differentiation of cells leading to a cancer stem cell phenotype and more aggressive tumors^63,64^. Our *MEN1* mutant tumors showed overexpression of *EZH2*, increased nuclear and cytoplasmic localization of EZH2, and decreased expression and membrane localization of E-cadherin. Interestingly, *EZH2* mutations in *MEN1* wild-type PanNETs showed an increase in H3K27me3 staining suggesting that an increase in activity of the polycomb repressive complex may play a role in PanNET progression and de-differentiation.

One interesting result was the ITGA2 staining in our PanNET tissues. Tissue expression of ITGA2 from the Human Protein Atlas exhibited staining in the nucleoplasm in immortalized HaCaT and U-2 OS cancer cells. *ITGA2* has also been shown to be upregulated in colon cancer^65^. We observed a statistically significant increase in nuclear ITGA2 staining in our *MEN1* mutant PanNETs perhaps due to the increase in *ITGA2* message present in these tumors. ITGA2 has also been shown to be a marker of stemness in prostate, uterine, and breast progenitor cells^66-68^. Therefore, the increase in ITGA2 staining in our *MEN1* mutant PanNETs may suggest that these tumors are more de-differentiated than *MEN1* wild-type tumors.

Angiogenesis is the development of new blood vessels and plays a critical role in the growth of cancer. Through the release of angiogenic growth factors, tumors induce angiogenesis resulting in new blood vessels within the tumor^69^. These newly-formed vessels then provide the tumor with oxygen and nutrients in addition to supplementary entry points for metastatic cells to enter the bloodstream to travel to distant sites within the body. Recently, several groups have shown the presence of cancer-derived endothelial cells suggesting that cancer stem-like cells are capable of undergoing differentiation into endothelial cells. Ricci-Vitiani et al. showed that a variable number (range 20-90%) of endothelial cells in glioblastoma carry the same genetic alterations as the primary tumor cells indicating a neoplastic origin of the vascular endothelium^70^. In vitro culture of a glioblastoma stem cell population in endothelial cell culture conditions generated progeny with phenotypic and functional endothelial cell characteristics^70^. Additionally, it has recently been shown that overexpression of Twist1 leads to endothelial differentiation of head and neck cancer^71^. Therefore, our results suggest that the increase in angiogenesis seen in our *MEN1*/*FGFR3* mutant PanNETs may be due to increased levels of de-differentiated cells in these tumors; however, more research is necessary to determine the mechanism behind increased angiogenesis in PanNETs.

Clinical management of patients with PanNETs is largely empirically based with no clear standard of how and when to treat the patient. Currently, surgical resection is the gold standard; however, for patients with unresectable or metastatic disease, treatment options are limited. In our set of PanNETs, we discovered mutations in several genes for which FDA approved therapies exist. *MTOR* or *TSC2* was mutated in 2/6 (33%) of our patients. The RADIANT-1-4 trials showed increased progression-free survival of patients with metastatic nonfunctional neuroendocrine tumors^72^. Everolimus was generally well tolerated during the trials; however, some patients required dose modification to complete treatment. Tyrosine kinases were mutated in 5/6 (83%) of our patients which included *SRC, FGFR3, AXL*, and *ALK*. Patients harboring mutations in these genes may benefit from any of the FDA approved oral tyrosine kinase inhibitors which have proved efficacious in multiple other cancers. Patients with *SRC* mutations may benefit from treatment with Dasatinib^73^ while patients with *AXL* mutations could potentially be treated with Cabozantinib^74^ and patients with *ALK* mutations given Crizotinib^75^. Recently, the FDA approved pan-FGFR inhibitor Erdafitinib for the treatment of locally advanced or metastatic disease in patients with bladder cancer harboring *FGFR2* or *FGFR3* mutations^76^. Interestingly, it has recently been shown that mutations in *KDM6A* amplifies polycomb repressive complex 2-regulated transcriptional repression which increased sensitivity to EZH2 inhibition^77,78^. Presently, the FDA has announced a fast-track investigation into a novel EZH2 inhibitor, Tazemetostat, for treatment of epitheloid sarcomas; though, further investigation on the efficacy of this inhibitor in cancers containing mutations in *KDM6A* is necessary.

Genomic changes responsible for the initiation and progression of pancreatic neuroendocrine tumors is poorly understood. In this study, we identified several signaling pathways potentially deregulated downstream of mutant *MEN1* along with additional novel therapeutic targets including *FGFR3* and *EZH2*. Currently, cellular models for sporadic, non-functional pancreatic neuroendocrine tumors do not exist. Development of mouse models for *in vivo* work or patient-derived organoid culture for *in vitro* work are necessary to fully characterize the mechanisms behind signaling downstream of mutant *MEN1* and mutant *FGFR3*. Additionally, determining the efficacy of currently available targeted chemotherapies is necessary to develop a better, more standardized treatment plan to improve treatment response and overall survival for patients with pancreatic neuroendocrine tumors.

## Acknowledgements

We would like to thank William Montfort for discussions involving potential functionality of the *FGFR3* mutations.

